# Developmental outflow tract abnormalities of Jag1-deficient mice are associated with abnormal ventricular activation and desynchronized contraction

**DOI:** 10.1101/2025.08.08.669322

**Authors:** Kristýna Neffeová, Eva Nekvindová, Veronika Olejníčková, Hana Kolesová

## Abstract

The Notch signaling pathway is an evolutionarily conserved intercellular communication mechanism essential for mammalian embryonic development. Mutations in the human *Jagged1* (*Jag1*) gene, which encodes a ligand of the Notch receptor, cause Alagille syndrome—an autosomal dominant disorder frequently associated with congenital heart diseases (CHDs) such as Tetralogy of Fallot.

To investigate the role of *Jag1* in cardiac development, we generated *Jag1^flox/flox^ Islet1-cre+* mice with a conditional deletion of the *Jag1* gene in the cardiac outflow tract. Mice carrying this targeted deletion exhibited severe cardiac malformations characteristic of Tetralogy of Fallot. The predominant defect observed was a double outlet right ventricle (DORV), in which both the aorta and pulmonary trunk arise from the right ventricle. This abnormality was consistently associated with a ventricular septal defect (VSD), present in 100% of homozygous mutants. Additional defects included abnormalities in the morphology of atrioventricular and semilunar valves, most commonly presenting as myxomatous mitral valves or altered leaflet numbers.

Since *Islet1* is also expressed in the sinoatrial and atrioventricular nodes, we employed optical mapping to visualize the cardiac conduction system. Analysis of E14.5 embryos and adult mice revealed altered activation patterns. While control hearts displayed a mature apex-to-base activation with conduction through both bundle branches, mutant embryos exhibited abnormal activation initiating exclusively from the left ventricle, indicating right bundle branch block (RBBB). In adult heterozygotes, electrical activation was asynchronous and often originated from ectopic sites, particularly in the posterior ventricular wall, deviating from the normal apical initiation observed in controls.

For functional analysis, we employed high-resolution ultrasound (Vevo imaging) on adult heterozygotes, as homozygotes did not survive postnatally. Most hemodynamic parameters showed no significant changes, suggesting early compensatory mechanisms to maintain cardiac output under compromised conditions. However, speckle-tracking strain analysis identified localized contractile defects and mechanical dyssynchrony, particularly affecting the anterior wall.

In summary, our study demonstrates that conditional deletion of *Jag1* leads to both morphological and electrophysiological abnormalities in the heart. These defects were evident in both homozygous and heterozygous embryos, with adult heterozygotes displaying persistent electrophysiological and mechanical alterations. The data suggest that disruption of *Jag1*-dependent signaling contributes to the pathogenesis of Tetralogy of Fallot and affects both the structural development and electrical function of the heart.

## Introduction

### Congenital heart diseases (CHDS)

CHDS affects 9.5 per 1000 live births^1^, constitutes a major fraction of clinically significant birth defects and represents an essential component of pediatric and adult cardiovascular diseases. Although there have been tremendous advances in diagnosing and treating CHDS, the knowledge of their causes is limited.

CHDS is a defect in the structure of the heart or great vessels. Most congenital heart defects are not associated with other organ diseases. CHDS can be divided into several categories: **hypoplasia** (underdeveloped right or left ventricle), **outflow tract defects** – mispositioning of aorta and pulmonary trunk, **obstructive defects** – narrowed or blocked outflow tract vessels and/or valves (pulmonary atresia, aortic atresia), **septal defects** (ventricular or atrial defect of the septa) and others. A common severe form of CHDS is the **Tetralogy of Fallot**, where four heart malformations are present (pulmonary atresia, ventricular septal defect, right ventricular hypertrophy, aorta connected to both ventricles^2^.

### Jagged1 (Jag1)

Jagged 1 (Jag1), a ligand for the Notch receptors, was identified as the gene responsible for Alagille syndrome^3^. Symptoms of this inherited disease may include Tetralogy of Fallot ^4^. Mice homozygous for a targeted null mutation of the Jag1 gene die at ED10 due to vascular defects ^5^.

Jag1 exhibits a temporal and spatial expression pattern during embryonic development. Jag1 expression is subsequently observed in the pharyngeal arches 1, 2, 3, 4, and 6, as well as in the heart, eye, and intersomitic spaces by ED9.5. Transmural Jag1 expression was detected in the atria and lower level of Jag1 expression is present in the ventricles at ED10.5. Robust Jag1 expression is detected in the descending aorta, the aortic arch arteries and the neural tube and the endothelial cells of the endocardial cushions within the atrioventricular canal ^6^. Jag1 was identified in sinus venosus endocardium and in endocardium that extended into the right atrium at ED11.5. Jag1 and Dll4 were expressed in the developing coronary arteries endothelium at ED13.5, in subsequent stages Jag1 expression is limited only to arterial endothelium ^7^. Jag1 is also expressed in a ductus arteriosus explaining patent ductus arteriosus in some patients with Alagille syndrome caused by Jag1 mutation ^6,8^. Besides arterial endothelium Jag1 is expressed in arterial perivascular smooth muscle cells^6,9^. In the ventricle, expression is more restricted to the endocardium and epicardium and Jag1 is not present in the myocardium ^6^. Information about the importance of Jag1-Notch signaling for myocardial and coronary development and its signaling in CHDS has been summarized in our review ^10^ (Neffeová, et al., 2022).

### Islet1

In this study we took advantage of silencing Jag1 in outflow tract region using *Jag1^flox/flox^ Islet1-cre+* mouse line. Islet1 is a marker of cardiac progenitor cells derived from secondary heart field and subpopulation of neural crest cells ^11^. Expression of the Islet1 can be firstly detected at stage ED7 (at the cardiac crescent stage) ^12,13^ and gradually decreases with the differentiation of cardiac progenitor cells ^13,14^. Those Islet1 positive cardiac progenitor cells differentiate into the three main cell types in the heart, cardiomyocytes, endothelial cells, and smooth muscle cells. Most cells within the outflow tract, right ventricle, ventricular septum and right atria, and a portion of cells within left ventricle and left atria derived from Islet1 expressing progenitors ^13^. Therefore, the hearts of mice missing Islet1 are completely lack the outflow tract, the right ventricle, and many of the atria mutant hearts and do not undergo looping^12^. Mice with homozygous deletion of Islet1 die at ED9.5-ED10.5 (with abnormalities in the organization of vascular endothelium) ^15^. Islet1 is also expressed in the wall of main stems of coronary arteries as well as in cardiac nerves ^13^

Here we report severe CHDS of Double outlet right ventricle (DORV) with VSD present in embryonic development of *Jag1^flox/flox^ Islet1-cre+* mouse line, which is so severe that the majority of models possessing this CHDS die perinatally. However, mainly *Jag1^flox/WT^ Islet1-cre+* mice surviving up to an adulthood and presents milder cardiac defect as abnormal heart activation patterns and irregularities in ventricular contraction and volume of the ventricles. Variability in cardiac phenotype of *Jag1^flox/flox^ Islet1-cre+* mouse line presented in embryos (DORV, VSD) is postnatally limited only to slightly abnormal morphology and function of the ventricles.

## Results

To elucidate and link the morphological and physiological changes associated with the development of severe congenital heart disease (CHDS) we took advantage of mouse model of conditional knockout of Jag1 (CKO Jag1). The conditional deletion of Jag1 under the promotor of Islet1, which is expressed in second heart field as well as in neural crest cells ^16^ results in severe malformations of outflow tract ^17^ - *Jag1^flox/flox^ Islet1^cre+^.* However the primary effect on the heart is double outlet right ventricle (DORV), this pathology also leads to the formation of a ventricular septal defect (VSD). Furthermore, maldevelopment of the outflow part of the heart is associated with abnormalities of both the outflow tract valves as well as in atrio-ventricular valves. In addition to structural defects, we also observed altered electrical activity and a decline in cardiac mechanical function, further underscoring the complex physiological consequences of Jag1 deficiency during heart development.

### Expression of Islet1

To visualize where Jag1 signaling was silenced in the heart, we use an advantage of reporter mouse ROSA-TdTomato, which we crossbred with the Islet1-cre mouse line. A red fluorescent signal of TdTomato in whole mounts of adult hearts was observed especially in the outflow tract (aorta and pulmonary trunk) (Fig 1. B), both atria and mosaic in both ventricles (Fig 1. A).

**Fig. 1.**
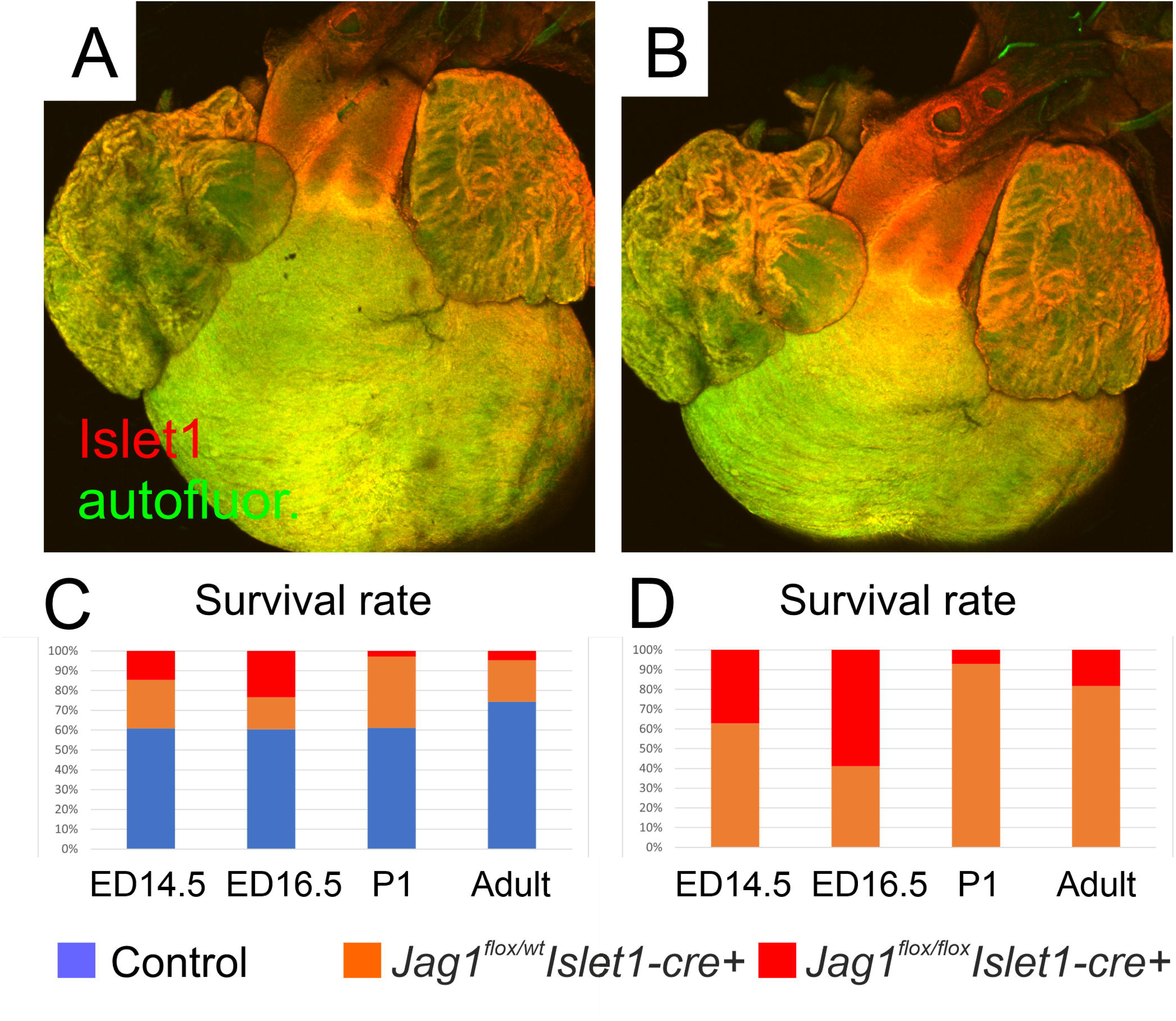
Expression of Islet1 in aorta, pulmonary trunk, outflow part of right ventricle, both atria and mosaic in the ventricles. Adult mouse heart from superior anterior aspect (A) and superior aspect (B). Survival rate of CKO Jag1 showing perinatal mortality of Jag1^flox/flox^ *Islet1-cre+* (C) and survival mainly of Jag1^flox/WT^ *Islet1-cre+* (D).

### Morphological changes in Jag1 CKO embryo (ED14.5 and ED16.5) – variability and Double outlet right ventricle (DORV)

Analysis of the late-gestation (ED14.5) hearts from *Jag1^flox/flox^ Islet1-cre+* and *Jag1^flox/WT^ Islet1-cre+* embryos revealed a number of severe cardiac abnormalities. The most prominent defect was DORV, where pulmonary trunk, as well as aorta, are both connected to the right ventricle (Fig. 2 A, D, G). The left ventricle is therefore drained via persisting VSD to the aorta (Fig 2. E, H). Embryos also exhibited malformations and thickening of the atrioventricular valves. (Fig 2. F, I). DORV was observed in 14% of *Jag1^flox/WT^ Islet1-cre+* and 57% of *Jag1^flox/flox^ Islet1-cre+* hearts.

**Fig. 2.**
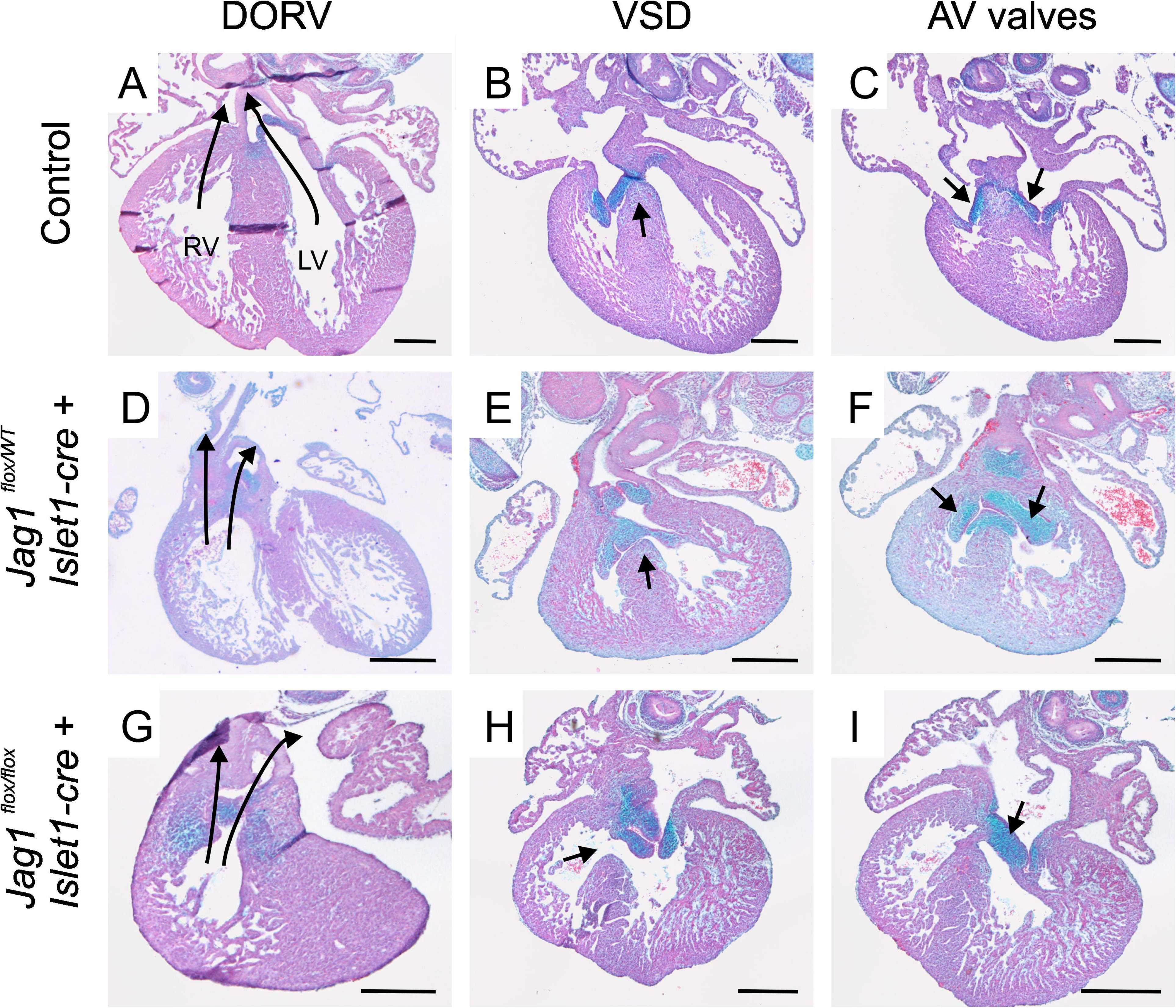
Representative morphological changes observed in at ED14.5 Jag1 CKO embryos with Double outlet right ventricle (DORV) and Ventricular septal defect (VSD). Normal development of ED14.5 with no double outlet (A), with fully closed interventricular septum (B) and with normal AV valves (C). Jag1^flox/WT^ Islet1-cre+ with double outlet right ventricle (D), presence of membranous VSD (E) and thickening and malformations of AV valves (F). Jag1^flox/flox,^ Islet1-cre+ with double outlet right ventricle (G), presence of VSD (H) and AV valves thickening and malformations (I). Right and left ventricle are placed similarly as on the (A). Scale is 200 um.

In DORV hearts, blood from the left ventricle is not able to continue directly to the aorta, which is connected to the right ventricle. This leads to non-closure of ventricular septum and to VSD. VSD enables the blood to continue from the left ventricle to the aorta. In our model none of the control (*Jag1^flox/flox^ Islet1-cre-*) littermates exhibit DORV and VSD, while in *Jag1^flox/WT^ Islet1-cre+* 2 from 7 analyzed embryonic hearts have DORV with VSD and all 7 analyzed embryos of *Jag1^flox/flox^ Islet1-cre+* of ED14.5 exhibit DORV with VSD.

The Jag1 CKO mouse model exhibits a spectrum of defects. We found that 71% of embryonic hearts have pulmonary atresia or atresia of the whole outflow tract. Increased resistance in pulmonary trunk resulted in thickening and hypertrophy of the right ventricle (14%).

Additional malformations observed in Jag1 CKO hearts included atrio-ventricular valve thickening and structural abnormalities (Fig. 2 C, F, I) ^18^. 51% of the hearts in our analysis showed distinct thickening of the atrio-ventricular valves already remarkable at the embryonic development. In some cases (30**%**) the fibrous ring and cardiac skeleton, to which the valves are attached, were affected and malformed in Jag1 CKO. In some cases (25%) valve leaflets also showed thickening in their distal parts.

Besides the gross malformations we observed that Jag1 CKO hearts show also fewer profound anomalies as developmental delay, delayed myocardial compaction and increased ventricular trabeculation.

The Jag1 CKO hearts have reduced functional fitness compared to controls, as only 8 out of 12 analyzed hearts of *Jag1^flox/flox^ Islet1-cre+* were spontaneously beating, in contrast to controls and *Jag1^flox/WT^ Islet1-cre+,* where only 2 out of 15 hearts were non-beating during optical mapping analysis (Fig. 3 G).

**Fig. 3.**
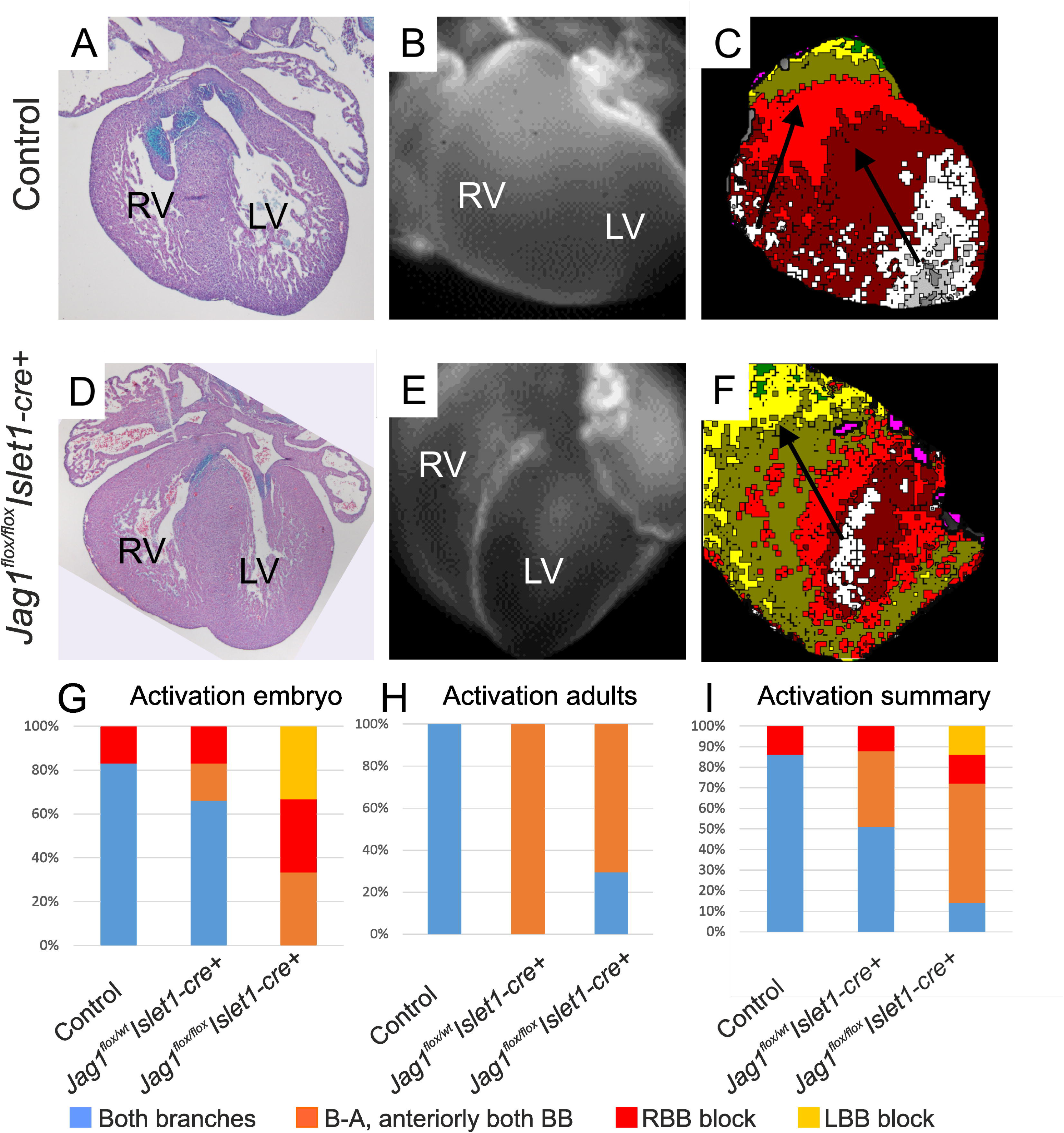
Activation patterns of embryonic hearts. Control heart (A) with normal activation pattern from both bundle branches (RBB and LBB) (B, C). Jag1^flox/flox^ Islet1-cre+ heart (D) with activation from LBB (RBB block) (E, F). Activation patterns of embryonic hearts (G-graph) where Jag1^flox/flox^ Islet1-cre+ with abnormal activation pattern. Activation patterns in adult hearts (H-graph) - disrupted activation of posterior heart aspect (B-A: base to apex). Summary of activation patterns with normal activation via both branches, abnormal activation from base to apex (B-A) but with activation via both branches and activation only with right or left bundle branch (RBB or LBB respectively).

DORV, which is predominantly observed in *Jag1^flox/flox^ Islet1-cre+*, represents a severe form of CHDS. Consequently, our model exhibits significant perinatal mortality, leading to a shift in the genotypic ratio of *Jag1^flox/flox^ Islet1-cre+* as the development proceeds. Overall, in our breeding cohorts at ED14.5 and ED16.5, we observed 60% controls, 25% of *Jag1^flox/WT^ Islet1- cre+* and only 15% of *Jag1^flox/flox^ Islet1-cre+* embryos, suggesting abortions of homozygotes. However, the ratio is significantly changing postnatally, with increased perinatal mortality especially of *Jag1^flox/flox^ Islet1-cre+*. The transition from fetal to postnatal (pulmonary) circulation is critically impaired in presence of severe CHDS such as DORV with VSD This is reflected in the ratio at postnatal day 1 (P1) mice where we still observed 60% controls and a notable proportion of Jag1^flox/WT^ Islet1-cre+ mice, which typically display a mild phenotype. In contrast, *Jag1^flox/flox^ Islet1-cre+* pups were represented by only a few percent, and those that did survive to P1 showed only mild or no detectable malformations.

Importantly, none of the *Jag1^flox/flox^ Islet1-cre+* animals exhibiting DORV or VSD (Fig. 1C, D) survived to adulthood, suggesting a strong negative selection against severe cardiac phenotypes postnatally.

### Ventricular activation pattern in embryonic heart

In order to link morphological abnormalities (DORV, VSD) to cardiac electrophysiology, we used optical mapping to analyzed ventricular activation patterns. Control heart exhibit mature activation pattern via both bundle branches by ED14.5. Only one control heart exhibit right bundle branch block (RBBB). On the other hand, both *Jag1^flox/WT^ Islet1-cre+* and *Jag1^flox/flox^ Islet1-cre+* embryos revealed severe alteration of activation patterns. Normal ventricular activation was not detected in any of *Jag1^flox/flow^ Islet1-cre+* hearts. The conduction abnormalities included immature base to apex activation in the posterior ventricular aspect, RBBB (Fig. 3 D-F) or left bundle branch block (LBBB). *Jag1^flox/WT^ Islet1-cre+* hearts showed normal ventricular activation in 66 % of the hearts (Fig. 3 G-I).

### Morphological changes in P1 and adult hearts

Hearts of *Jag1^flox/WT^ Islet1-cre+* mice surviving to to P1 exhibited morphology similar to control hearts, with only small percentages of animals showing DORV with VSD (Fig. 4 D, E), while the remainder appeared indistinguishable from controls (data not shown). In adult Jag1 CKO none of them exhibit DORV with VSD. Overall, heart gross morphology of Jag1 CKO (*Jag1^flox/WT^ Islet1-cre+*) was comparable to controls (Fig. 4 C, F) with no profound structural defect. This could imply that the animals with the most severe heart defect died perinatally and are not included in the further analyses. On the other hand, as the hearts had in their history occurrence of abnormalities of outflow tract area, the morphological analysis of the outflow tract area (base of the heart) revealed certain anomalies. These includes extra tissue and incoherencies in the smooth muscle wall of aorta and pulmonary trunk (Fig. 5 K, L). We also found improper anulus fibrosus formation in atrioventricular connection with myocardial continuity, which could results in altered heart activation and ventricular pre-excitation (Fig. 5 O, P).

**Fig. 4.**
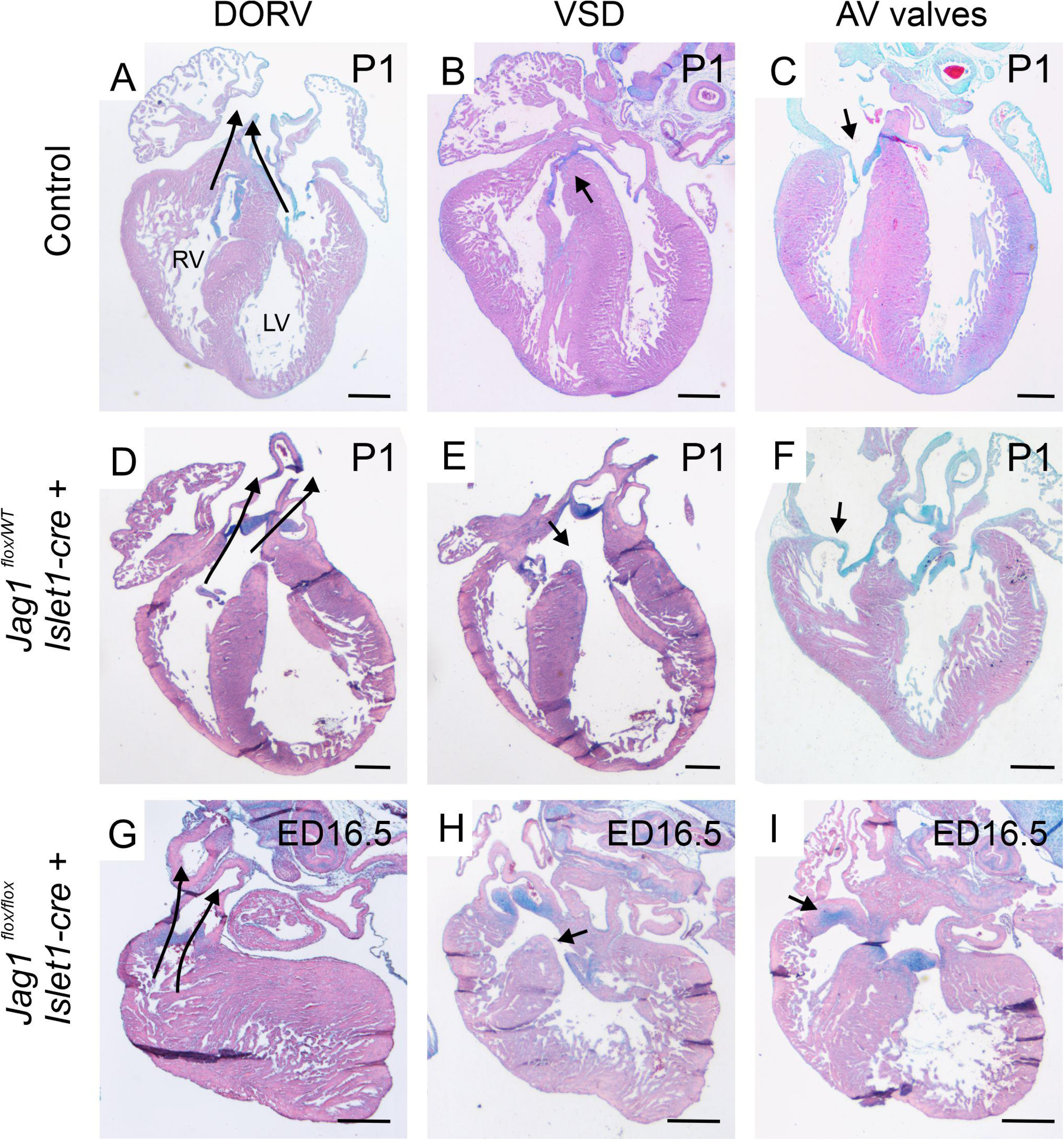
Morphological abnormalities in ED16.5 and P1. Double outlet right ventricle (DORV) on Jag1 CKO. Normal development P1 without double outlet (A), with fully closed interventricular septum (B) and with normal AV valves (C). Jag1^flox/WT^ Islet1-cre+ with double outlet right ventricle (D), presence of membranous VSD (E), AV valves are normal (F). Jag1^flox/flox^ Islet1-cre+ with double outlet right ventricle (DORV) (G), presence of VSD (H) and AV valves thickening and malformations (I). Right and left ventricle are placed similarly as on the (A). Scale is 500 um.

**Fig. 5.**
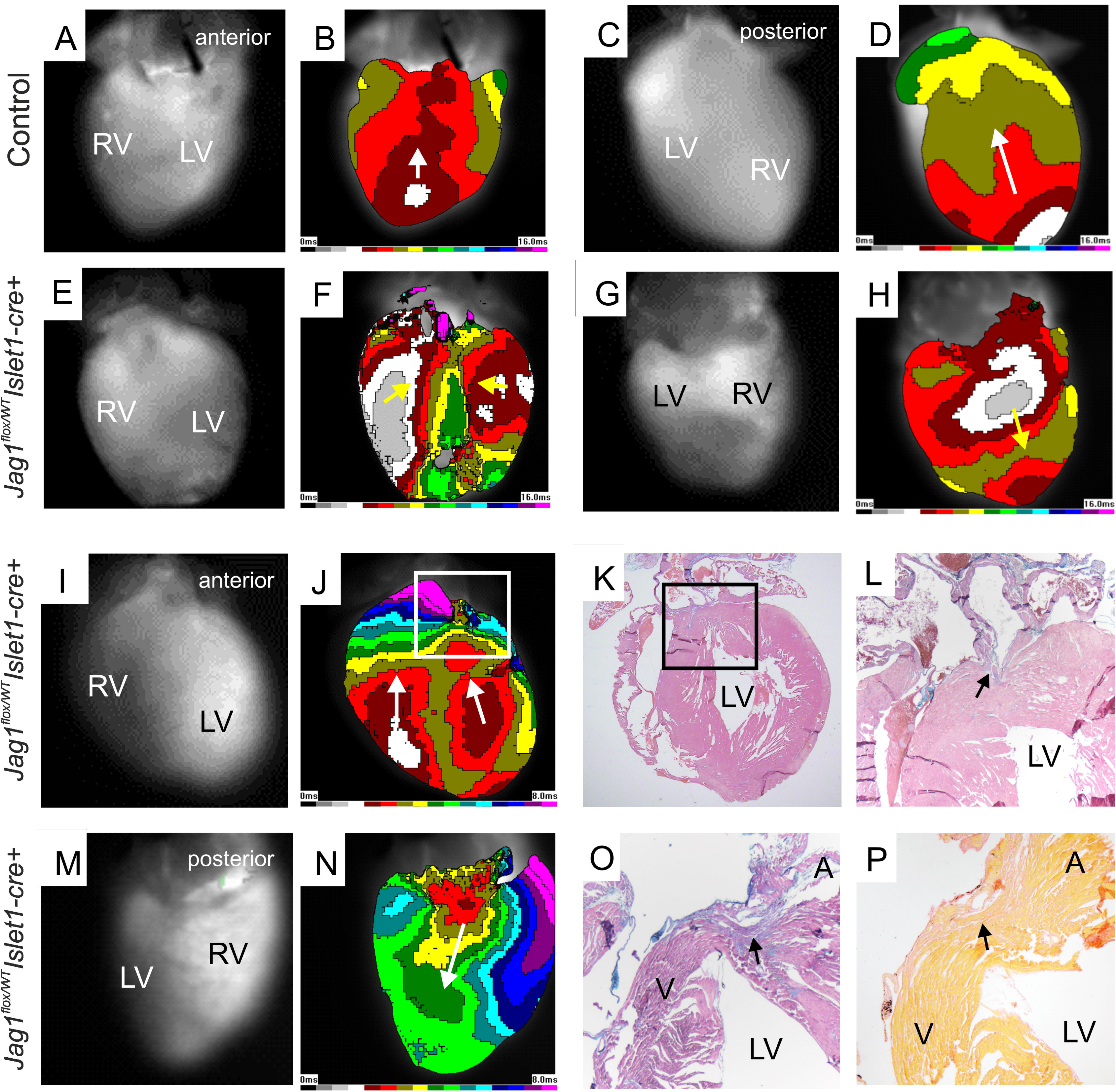
Activation patterns of adult hearts. The adult control heart is activated from the apex and the signal is continuing to the base (A, B, C, D). Jag1^flox/WT^ Islet1-cre+ is activated by LBB and RBB in the middle of ventricular length anteriorly (E, F) and posteriorly from middle of the ventricle (G, H). Irregularities in Jag1^flox/WT^ Islet1-cre+ activation of the OFT (I, J), discontinuity of myocardium in OTF visualized by histology-ABHE (K, L). Posteriorly Jag1^flox/WT^ Islet1-cre+ hearts are activated from base to apex (M, N), lack of fibrous insulation between atria and ventricles, myocardial continuity (P, arrows). Histology-ABHE (O), picrosirius red staining (P). LV-left ventricle, RV-right ventricle.

### Ventricular activation pattern in adult hearts

Electrical activation of adult *Jag1^flox/WT^ Islet1-cre+* hearts showed differences from normal pattern. As we seen on control mouse hearts, ventricles are normally activated through both bundle branches. This path of impulse spreading is resulting in apex-first activation pattern revealing in both anterior and posterior aspect of the ventricle (Fig 5. A-D). However, similarly as in embryonic Jag1 CKO, we observed abnormal activation in *Jag1^flox/WT^ Islet1-cre+* hearts, where activation signal spread in opposite direction from the base of the ventricle and continue directly to the posterior aspect of the ventricle. (Fig 5. G, H, M, N). A milder phenotype was observed in other hearts, where the overall direction of activation was preserved, but the initial activation site was located laterally in both ventricles, with irregular propagation patterns ((Fig 5. E, F). Another observed variation involved abnormal conduction in the outflow tract region (Fig 5. I, J), which can be affected by loss of Jag1 function in outflow tract region (under the promotor of Islet1).

Histological analysis of the affected basal region of the heart using ABHE staining (Alcian blue for connective tissue) and picrosirius red (for collagen fibers) revealed disruptions in fibrous insulation between the atria and ventricles and areas of persistent myocardial continuity (Fig 5. O, P). Which correlates with improper activation from the base of the heart, where activation signal from the atria can continue directly and pathologicaly to the ventricle. Which is resulting in abnormal ventricular activation pattern from the base to apex (Fig. 5. G, H, M, N) ^19^. Similarly, irregular action potential spreading in the outflow part of the heart (Fig 5. I, J), where by histological analysis we found extra non-fibrous tissue in the area of annulus fibrosus, which can be responsible for the irregular activation (Fig 5. K, L).

Jag1 CKO mouse adult models are activated at the anterior aspect of ventricular wall from both bundle branches (Fig 5. E, F, I, J), even with some irregularities. However posteriorly activation pattern is more affected by loss of Jag1, and ventricular activation happens from base to apex (Fig 5. G, H, M, N).

### Physiological changes in adult hearts

To determine the effect of morphological abnormalities on heart function, we performed echocardiographic measurements using VEVO high-frequency ultrasound imaging. (Fig 6 and 7). Physiological parameters were assessed from M-mode short-axis view images. We analyzed the hearts of adult mice from the control group and compared them to those of the *Jag1^flox/WT^ Islet1-cre+* group. Our analysis revealed a comparable range of heart rates in both groups, as well as unchanged cardiac output, as depicted in Fig. 7, graphs A and B. However, we found a significant difference in fractional shortening (34.8 ± 2.2 in controls compared to 38.6 ± 2.0) (Fig. 7, graph C) and ejection fraction (64.5 ± 3.0 compared to 69.6 ± 2.6) (Fig. 7, graph D), both of which were higher in the *Jag1^flox/WT^ Islet1-cre+* group. Additionally, a significant difference was observed in left ventricular diameter during systole (DiameterS) (Fig. 7, graph E) (2.6 ± 0.1 in controls compared to 2.3 ± 0.2) and end-systolic volume (ESV) (Fig. 7, graph G) (24.5 ± 3.0 in controls compared to 17.9 ± 4.5), both of which were reduced in *Jag1^flox/WT^ Islet1-cre+* mice. However, these factors remained unchanged during diastole (Fig. 7, graph F, H). Specifically, the diameter of the left ventricle during diastole (DiameterD) (Fig. 7, graph F) was 4.0 ± 0.2 in controls and 3.7 ± 0.3 in *Jag1^flox/WT^ Islet1-cre+*, and end-diastolic volume (EDV) (Fig. 7, graph H) was 68.8 ± 7.2 compared to 58.0 ± 10.3.

**Fig. 6.**
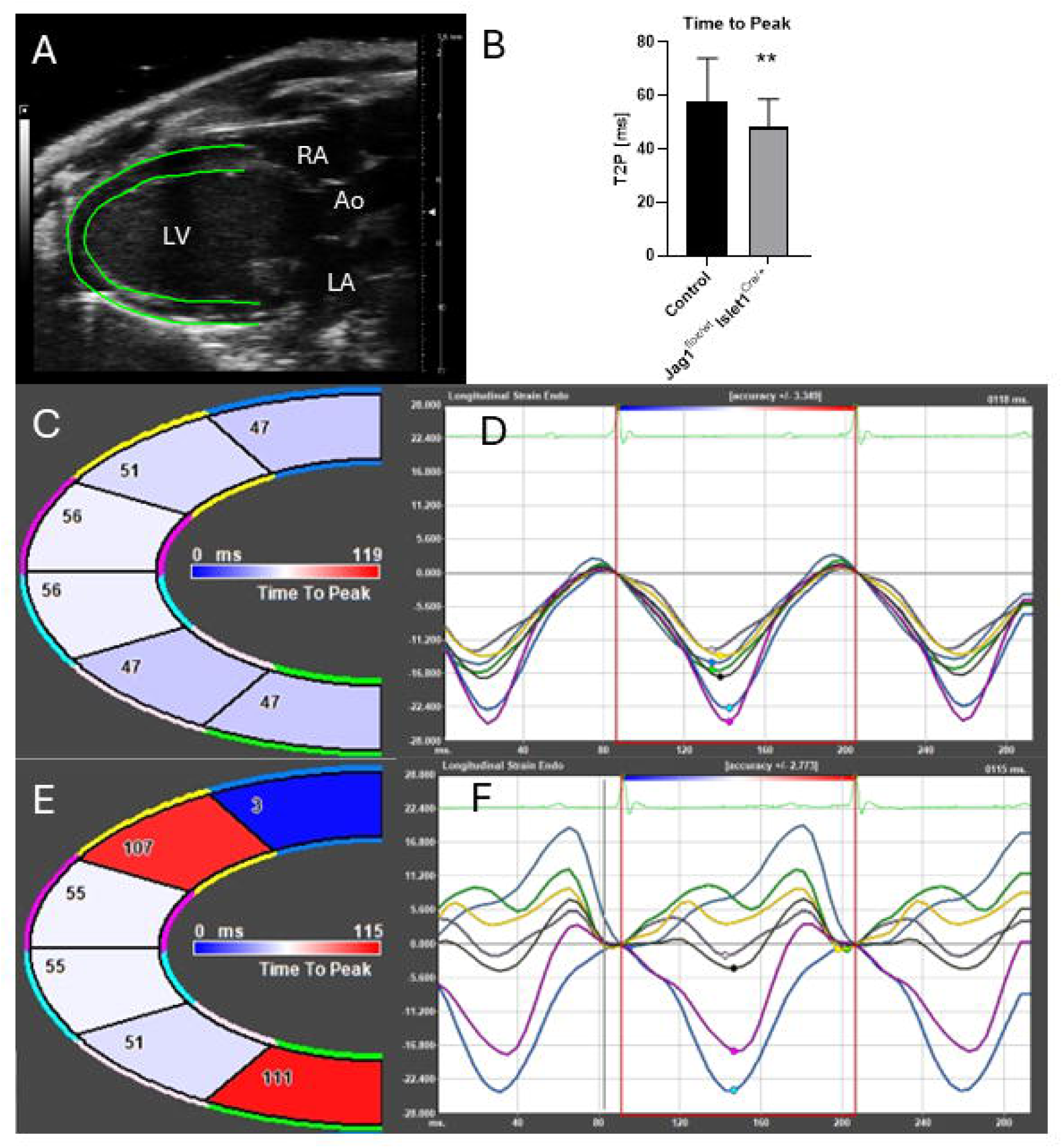
Echocardiography analysis of adult hearts. Left ventricle (LV) strain analysis was conducted using B-Mode images acquired from the parasternal long axis view, as illustrated in panel A. Speckle tracking was utilized to trace the endocardial and epicardial borders, depicted as green lines in panel A, and separately for each LV segment. LV segments are delineated in panels C and E, with the time to peak strain (T2P) indicated for each segment according to the color key. The colors of the segment edges correspond to the colors of the lines in panels D and F, representing the temporal progression through the cardiac cycle. Panels C and D represent the analysis results of a control mouse, while panels E and F depict Jag1^flox/WT^ Islet1-cre+. In Jag1^flox/WT^ Islet1-cre+ (E, F), desynchrony indicated by varying T2P compared to controls is evident, it also manifested as asymmetry among curves in panel F. Graph B quantifies strain as the time from baseline to peak strain (T2P), illustrating individual values for epicardial segments in longitudinal strain in both control and Jag1^flox/WT^ Islet1-cre+. Overall, a lower T2P of longitudinal strain in the epicardium was observed in the Jag1^flox/WT^ Islet1-cre+ group compared to controls.

**Fig. 7.**
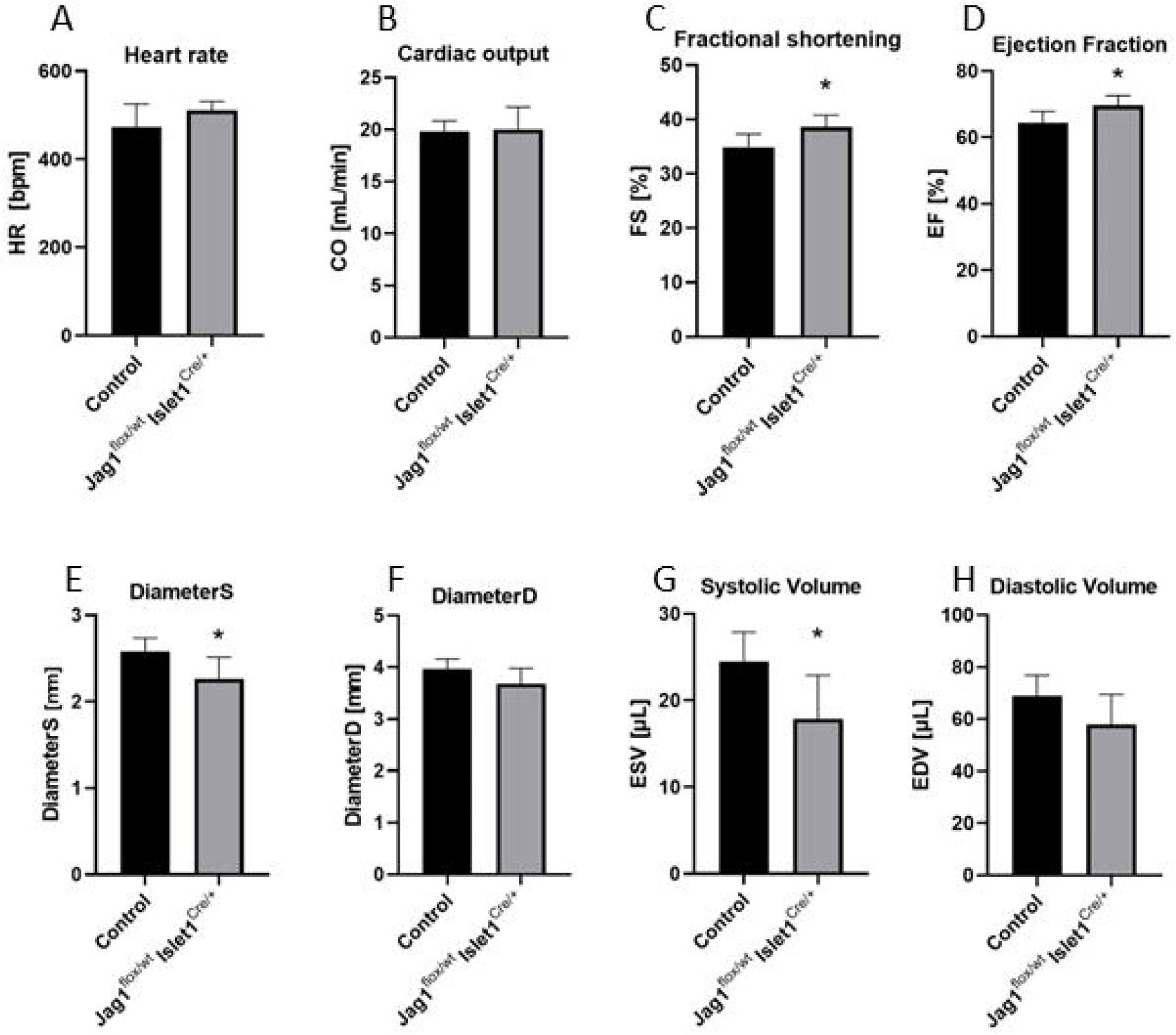
Echocardiographic analysis of short axis, M-mode. Echocardiography show trend in increase of the heart rate and cardiac output. Fractional shortening and ejection fraction is significantly increased in Jag1^flox/WT^ Islet1-cre+. Systolic diameter (DiameterS) is significantly decreased, however diastolic diameter (DiameterD) is non significantly decreases in Jag1^flox/WT^ Islet1-cre+. End ystolic volume (ESV) is significantly decreased in Jag1^flox/WT^ Islet1-cre+. Data suggest adaptive hypertrophy as a compensatory mechanism to improve myocardial contractility, to reduce wall stress and maintain cardiac output.

Strain analysis revealed contraction desynchrony in the left ventricle of *Jag1^flox/WT^ Islet1-cre+* mice, compared to the controls, as illustrated in Fig. 6, pictures C and E, where individual segments exhibited heterogenous values of time-to-peak (T2P) values, as well as asynchronous curves depicting ventricular contraction during the cardiac cycle (Fig. 7, pictures D and F). Differences were pronounced in both the longitudinal and radial directions of contraction.

Quantification of individual values of T2P revealed significantly reduced values in *Jag1^flox/WT^ Islet1-cre+* epicardial longitudinal strain (57.6 ± 16.1 in controls compared to 48.37 ± 10.2), as shown in Fig. 7, graph B, along with reduced contraction velocity of the anterior ventricular wall in *Jag1^flox/WT^ Islet1-cre+*. The posterior part of the free left ventricular wall appeared less affected compared to the anterior ventricular wall. Some segments exhibited significant variability in strain and displacement, suggesting regional mechanical inhomogeneity within the myocardium.

Taken together, adult mice exhibit ventricular contraction dyssynchrony, which likely results from impaired electrical conduction as detected by optical mapping. This conduction disturbance manifests functionally as mechanical dyssynchrony, confirmed by strain analysis using VEVO echocardiography.

## Discussion

*Jag1^flox/flox^ Islet1-cre+* mouse line exhibits Jag1 signalization silenced in outflow tract endothelium and smooth muscle cells of arterial vessels wall ^6,11^. Silencing of Jag1 function leads to severe morphological abnormalities in heart embryonic development, which in most severe form are manifested as Double outlet right ventricle (DORV), where aorta and pulmonary trunk are both connected to the right ventricle. As the left ventricle cannot be drained directly, the blood flow in embryonic development prevents the closure of the interventricular septum, which leads to ventricular septal defect (VSD). Due to the misplacement of the aorta, the entire region of the heart base is affected and malformed. These malformations include the malformation of valve leaflets, improper electrical insulation, which is manifested as persisting myocardial connections between the atria and ventricles. We propose that these morphological malformations of outflow tract area (base of the heart) cause the irregularities in cardiac electrical conduction, which is then manifested, in embryos as well as in adults, as abnormal activation patterns of both ventricles. From postnatal physiological analysis of our mice model, we see that abnormal heart activation leads to incoherencies of left ventricular contraction, manifested by various physiological parameters.

Our model of *Jag1^flox/flox^ Islet1-cre+* in embryonal development exhibit whole spectrum of defects, from almost normal phenotype, especially in *Jag1^flox/WT^ Islet1-cre+* ^T^to severe CHDS, including DORV and VSD. Analysis of heart activation also revealed that the most profound abnormalities in heart activation pattern (base to apex activation at the posterior aspect of the ventricle or bundle branches blocks) occurred in embryos with the most severe morphological defects (DORV, VSD). These findings demonstrate that morphological changes in the area of the heart base, have intensively influenced ventricular electrical activation and overall cardiac physiology, already during embryonic stages.

We assume that these morphological and physiological changes, caused by maldevelopment of outflow tract, are sufficiently severe to prevent adaptation to postnatal (pulmonary) circulation, ultimately resulting in perinatal lethality as supported by survival rate analysis.

Animals surviving to adulthood possess milder morphological phenotype, without DORV and, in most cases, no persistent VSD. This suggests that the most severely affected embryos do not survive the perinatal period, and that mild VSDs may close spontaneously after birth. However, these postnatal hearts morphologically resemble their littermate controls, their outflow tract area exhibit some minor morphological changes like improper electrical insulation between atria and ventricles or great vessels wall abnormalities. We think that these minor morphological changes are causing improper heart activation and, consequently, mechanical malfunction of ventricular contraction. These hearts exhibited incoherent ventricular contraction and reduced ventricular volumes.

Even though the adult hearts exhibit mild or almost no morphological phenotype, their history of improper heart development due to silenced Jag1 signaling results in important physiological abnormalities as altered heart activation pattern, incoherent ventricular contraction and affected ventricular volume. We suggest that these physiological abnormalities (remnants from embryonic development) ^20^ can cause in later life worsening of heart adaptive functions on the organism environment.

Echoardiographic results showed starting trend to cardiac hypertrophy (increased Fractional shortening and Ejection fraction as well as decreased Diameter S and End systolic volume), as an adaptive mechanism to improve cardiac contractility, reduce wall stress and maintain cardiac output^21^. Althought cardiac output remained stable, the significantly elevated Fractional shortening and Ejection fraction in the Jag 1 CKO group support the notion of early compensatory hypertrophy. Also, the presence of ventricular dyssynchrony may represent an early indicator of pathological remodeling^22^.

We observed lower diameter of left ventricle as well as end systolic volume indicating concentric hypertrophy. We hypothesize that maldevelopment of cardiac outflow tract could cause pressure overload resulting in pressure overload ventricles. Than concentric hypertrophy could develop as adaptive mechanism to improve myocardial contractility, reduce wall stress and maintain cardiac output as we seen in our data despite lower systolic diameter and volume. Compensatory response of heart also indicates elevated ejection fraction and fractional shortening. However, we hypothesize that in long term this adaptive mechanism could change to maladaptation and could lead to cardiac pathology. This would also correspond to dyssynchronical contraction of left ventricle revealed by speckle tracking analysis, because mechanical dyssynchrony is as an early marker of cardiomyopathic disease. Mainly because mechanical dyssynchrony progressed into global myocardial discoordination and decompensated cardiac pump function

However mild physiological heart abnormalities are challenging to detect due to methodological limitations, but they remain highly relevant, as they can have their origin in improper morphological embryonic development in different areas of the heart. This is manifested on *Jag1^flox/flox^ Islet1-cre+* model, where morphological changes in outflow tract resulted in improper ventricular activation and function.

## Material and Methods

### Experimental animal model

To study the role of Jag1 at specific Islet1 positive sites of the heart, we generated *Jag1^flox/flox^ Islet1-cre+* mice model, described earlier (High et al. 2009). For crossbreeding we used Jag1^tm2Grd^SjJ (RRID:IMSR_JAX:031272) and Islet1^tm^^1(cre)Sev^/J (RRID:IMSR_JAX:024242) mice on a C57BL/6J background (RRID:IMSR_JAX:000664). Islet1 ^tm1(cre)Sev^/J mice were kindly provided by Dr. Pavlínková (Laboratory of Molecular Pathogenetics, Institute of Biotechnology Czech Academy of Sciences, BIOCEV) ^23^. Jag1^flox/flox^ females were caged overnight with heterozygous *Jag1^flox/WT^ Islet1-cre+* males. In the morning, females were checked for vaginal plugs; the noon of the day of plug discovery was considered ED0.5. Mice without the Isl1-Cre allele were used as controls.

Animals were euthanized by cervical dislocation and collected at different ages – embryos ED14.5, ED16.5, neonatal P1 and adults. To visualize Islet1 location we crossbreed Islet1^tm1(cre)Sev^/J with Cre reporter mouse B6.Cg-Gt(ROSA)26So^rtm^^14(CAG-tdTomato)Hze^/J, provided by Dr. Mašek (Laboratory of Intercellular Communication, Faculty of Science, Charles University, Prague) ^24^.

Animals were housed in a controlled environment with 12h light/dark cycles with all applicable regulations with food and water ad libitum. All of the animal breeding and procedures were performed according to the approval by Center for Experimental Biomodels, First Faculty of Medicine, Charles University and Institute of Anatomy, First Faculty of Medicine, Charles University. This study was designed and reported in accordance with the ARRIVE 2.0 guidelines (Animal Research: Reporting of In Vivo Experiments).

Used genotypes are summarized in the Table of Genotypes 1. As controls we have analyzed Jag1^flox/flox^ mouse and Jag1^flox/flox^ Islet1-negative littermates without Cre recombinase.

**1.**
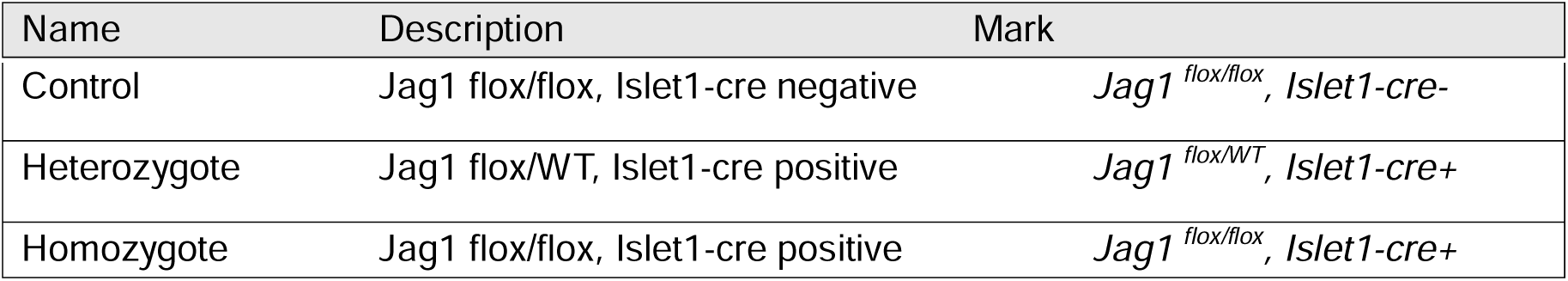
Table of the genotypes.

**1 Table of the genotypes**
| Name | Description | Mark |
| --- | --- | --- |
| Control | Jag1 flox/flox, Islet1-cre negative | *Jag1<sup>flox/flox</sup>, Islet1-cre-* |
| Heterozygote | Jag1 flox/WT, Islet1-cre positive | *Jag1<sup>flox/WT</sup>, Islet1-cre+* |
| Homozygote | Jag1 flox/flox, Islet1-cre positive | *Jag1<sup>flox/flox</sup>, Islet1-cre+* |

### Histology

All samples were perfused with cardioplegic solution, fixed overnight in 4% paraformaldehyde in phosphate buffer saline (PFA/PBS) and dehydrated stepwise with an ethanol-to-benzene series and embedded in Paraplast Plus (ROTH). The paraffin blocks were sectioned at a thickness of 8µm. Alcian blue-Hematoxylin-Eosin (ABHE) basic histological staining was used for morphological analysis. The staining times depended on the age of the animals.

Subsequently, Picrosirius red, collagen specific staining, was used for analysis of atrioventricular junctions. Adult heart sections were stained with 0,1% picrosirius red solution for 1 min and cardiomyocytes were then stained with saturated solution of picric acid for 4 minutes at room temperature. In the end, the sections were dehydrated in ascending series of ethanol, cleared in xylene, and mounted in Histokit medium. Sections were analyzed under transmitted light - microscope Olympus BX51 with CCD cameras Olympus DP71 and DP80.

### Optical mapping

To analyze the pattern of ventricular activation, optical mapping was performed at embryonic day ED14.5, ED16.5, and adult hearts. Time-pregnant females were sacrificed by cervical dislocation and embryos were kept in ice-cold oxygenated Tyrodés solution until individual dissection. After removal from extraembryonic membranes, embryos were sacrificed by rapid decapitation. Hearts were then isolated and stained by immersion in di-4-ANEPPS (ThermoFisher Scientific; 1.7 mM) for 12 minutes on ice ^25^. The excitation-contraction uncoupler +-blebbistatin (Sigma Aldrich; 0.1 µM) was added to prevent motion artifacts. Mapping of epicardial electrical activity was performed in spontaneously beating hearts immersed in warm (37°C Tyrodés solution (composition in mmol/l: NaCl 145 NaCl, 5.9 KCl, 1.1 CaCl2, 1.2 MgCl2, 11 glucose, 5 HEPES; pH = 7.4; gassed with 100% O2) using the Ultima L high-speed camera (SciMedia Ltd., Tokyo, Japan) fitted to a BX51 FS epifluorescence microscope (Olympus, Japan) and equipped with a 150 W Xe arc lamp (Cairn, UK) ^26^. Recordings were analyzed following established protocols using BV_ana software (SciMedia Ltd., Tokyo, Japan) ^27^. After mapping, hearts were fixed in 4% paraformaldehyde for 24 hours and used for subsequent analyses.

### Echocardiography - VEVO

Echocardiography was performed on animals aged between three to six months, following the protocol outlined in ^28^. Briefly, mice were anesthetized for 30 seconds using 5% isoflurane in an anesthesia box before being placed in a supine position with their paws embedded in electrode gel and secured with adhesive tape to enable the recording of breathing frequency and ECG signal (lead II). Core body temperature was monitored using a rectal probe and maintained at 37.5 ± 0.5°C using a heating plate and infrared heating lamp. Eye ointment was applied to prevent dehydration, while hair removal cream was applied on the animal’s chest for one minute. Echocardiography was performed under 1-2% isoflurane anesthesia delivered via a mask with oxygen at 1.5 L/min. The Vevo3100 preclinical imaging system (Fujifilm VisualSonics) with an MX550D transducer was utilized to capture B-mode and M-mode recordings in both long and short-axis views of the left ventricle. Data analysis was conducted using the Vevo®Lab software (Fujifilm VisualSonics), allowing for assessment of basic cardiac physiological parameters and speckle-based strain analysis, providing detailed insights into the synchronicity and motion of the left ventricle in radial and longitudinal directions.

For speckle tracking analysis, the left ventricle was divided into 6 segments. Analysis was conducted individually for each segment in terms of both longitudinal and radial movement for the epicardium and endocardium (Fig. 6). Strain was quantified by measuring the time taken to reach the highest point of the curve representing the strain pattern throughout the cardiac cycle, referred to as time to peak (T2P) in milliseconds.

## Competing interests

Authors declare no competing interests.

## Acknowledgement

We would like to thank for excellent technical assistance to Blanka Topinkova and Marketa Hemerova.

## Funding

Supported by Czech Science Foundation 201521 24-12330K, the project National Institute for Research of Metabolic and Cardiovascular Diseases (Program EXCELES, ID Project No. LX22NPO5104) - Funded by the European Union - Next Generation EU and Czech Health Research Council: NU21J-02-00039. Charles University institutional funding COOPERATIO (Charles University Cooperatio 207029 Cardiovascular science). CAPI and Czech-BioImaging project (MEYS, project number LM2023050 Czech-BioImaging).

## Data accessibility

Figures and data are publicly accessible at DOI: 10.5281/zenodo.16780062.

Preprint is accessible at BioRxiv.

Data are also deposited at Genome Phenome Archive.

## Author contribution statement

K.N.-data acquisition and figures preparation, manuscript preparation, E.N - data acquisition and figures preparation, V.O. - data acquisition and figures preparation. H.K – data acquisition, figures and manuscript preparation.

Zabrodska et al., 2024 DOI: <u>10.1152/ajpheart.00115.2024</u>

Olejnickova et al., 2022 DOI: <u>10.1002/dvdy.413</u>

Olejnickova et al., 2021 DOI: <u>10.3390/ijms22052475</u>

Kvasilova et al, 2020 DOI: <u>10.1242/jeb.229278</u>

## Notes

### Competing Interest Statement

The authors have declared no competing interest.

